# Coordination of two opposite flagella allows high-speed swimming and active turning of individual zoospores

**DOI:** 10.1101/2021.04.23.441092

**Authors:** Quang D. Tran, Eric Galiana, Philippe Thomen, Céline Cohen, François Orange, Fernando Peruani, Xavier Noblin

**Author notes:** Institut Jacques Monod (IJM), University of Paris, CNRS, UMR 7592, 75013 Paris, France.

## Abstract

*Phytophthora* species cause diseases in a large variety of plants and represent a serious agricultural threat, leading, every year, to multibillion dollar losses. Infection occurs when these biflagellated zoospores move across the soil at their characteristic high speed and reach the roots of a host plant. Despite the relevance of zoospore spreading in the epidemics of plant diseases, characteristics of individual swimming of zoospores have not been fully investigated. It remains unknown about the characteristics of two opposite beating flagella during translation and turning, and the roles of each flagellum on zoospore swimming. Here, combining experiments and modeling, we show how these two flagella contribute to generate thrust when beating together, and identify the mastigonemes-attached anterior flagellum as the main source of thrust. Furthermore, we find that turning involves a complex active process, in which the posterior flagellum temporarily stops, while the anterior flagellum keeps on beating and changes its pattern from sinusoidal waves to power and recovery strokes, similar to *Chlamydomonas*’s breaststroke, to reorient its body to a new direction. Our study is a fundamental step towards a better understanding of the spreading of plant pathogens’ motile forms, and shows that the motility pattern of these biflagellated zoospores represents a distinct eukaryotic version of the celebrated “run-and-tumble” motility class exhibited by peritrichous bacteria.

## INTRODUCTION

Life of swimming microorganisms in viscosity-dominant world has been of great interest in biophysics research. The problems on microbial locomotion of those tiny individual flagellated swimmers are still far to be fully understood. There have been a multitude of theoretical and experimental models of microswimmers that study the hydrodynamics of the individual and collective motions of those cells [1, 2]. These swimming cells can be categorized into two groups: eukaryotes (having nuclei) and prokaryotes (no nuclei). *Escherichia coli* is one of the most studied prokaryotic swimmers, which possesses a bundle of passive helical flagella controlled by a rotary motor attached to the cell body [3]. Eukaryotic microswimmers, such as green algae *Chlamydomonas* [4, 5] and spermatozoa [6, 7], have active *and flexible flagella along which molecular motors are distributed. Here, we introduce a new type of microswimmer, named Phytophthora* zoospores, which has two different flagella collaborating for unique swimming and turning mechanisms (Figure 1(A)).

**FIG. 1.**
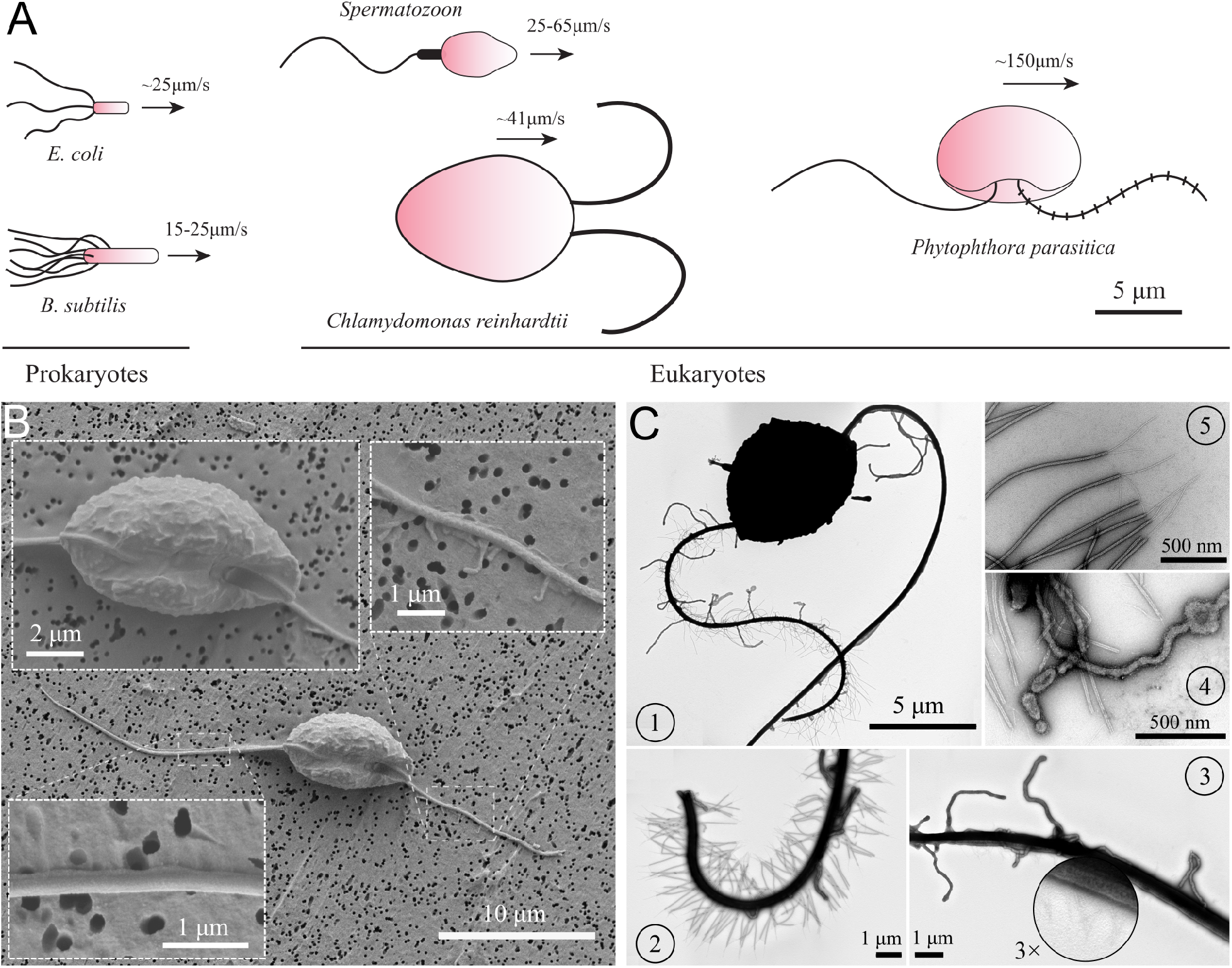
Characteristics of *P. parasitica* zoospore. (A) Swimming of zoospores in comparison with different prokaryotic and eukaryotic microswimmers. Black arrows indicate the swimming direction of the swimmers. (B) Scanning electron microscopy images of the zoospore. The insets show the enlarged images of the cell body and the two flagella. (C) Transmission electron microscopy images with negative staining. (1) Image of the zoospore showing the different structures of the two flagella. The anterior has multiple mastigonemes, while the posterior has a smooth straight structure. (2) Close zoom-in image of the anterior flagellum. It is noticed that there are two types of mastigonemes on this flagellum: one with straight tubular shape, the other with flimsy shape but longer and bigger in size. (3) Close zoom-in image of the posterior flagellum. There are plenty of thin and short hairs wrapping along the flagellum and several non-tubular mastigonemes appearing near the cell body. (4) The non-tubular mastigonemes. (5) The tubular mastigonemes with tiny hairs at the tip.

*Phytophthora* is a genus of eukaryotic and filamentous microorganisms. They are classified as oomycetes and grouped in the kingdom of the Stramenopiles with the heterokont algae (such as diatoms and brown algae) [8, 9]. A number of *Phytophthora* species are plant pathogens and cause tremendous damages to agro- and eco-systems [10, 11]. Nowadays, *Phytophthora* diseases are responsible for a big impact on economies with billions of dollars of damages each year and remain a threat to the food security worldwide [12–14]. The diseases are pervasive as they release swimming biflagellated spores called “zoospores” which initiates the spreading through water. These zoospores are able to achieve speed up to 250 µm s^−1^ [15] through thin water films, water droplets on leaves, or through pores within moist soils. To facilitate the spreading, their cell bodies store an amount of energy (mycolaminarin, lipid) allowing them to swim continuously for several hours [14]. In natural ecosystems and even more in agro-systems, putative host plants are usually close. This proximity makes the distance to find a plant relatively short and it is compatible to the time-ability of zoospores to swim. When the zoospores reach plant roots, they stop swimming and release their flagella to produce a primary cell wall and become germinative cysts which are able to penetrate into the host tissue. Then, they start a hyphal growth inside the infected plant. In this study, we investigate the telluric species *P. parasitica*, a polyphagous pathogen attacking a wide range of hosts such as tobacco, onion, tomato, ornamentals, cotton, pepper, citrus plants and forest ecosystems [16].

Previous studies have shown that during the spreading and approaching the host, zoospores can have complicated swimming patterns and behaviors as they experience multiple interactions with environmental signals, both physical, electrical and chemical, in soil and host-root surface [17]. Near the plant-root, zoospores can perceive various stimuli from the environment, such as ion exchange between soil particles and plant roots, the chemical gradients generated by root exudates, which activate cell responses. This results in coordinated behaviors of zoospores, allowing them to preferentially navigate to the water film at the interface between soil particles and plant roots. For instance, potassium, which is uptaken by roots in the soil, reduces zoospore swimming speed, causes immediate directional changes and also results in perpetual circle trajectories [15, 18]. Bassani *et al*. provide transcriptomic studies showing that potassium induces zoospore aggregation, which facilitates the advantages for zoospores to attack the host-root [19]. Experimental evidence has demonstrated that zoospore-zoospore interaction can lead to “pattern swimming”, a microbial bioconvection happened without the appearance of chemical or electrical signals [20, 21]. These findings urge for a better understanding of the swimming physics of the individual zoospores and how the combination of their two heterogeneous flagella results in those complex swimming behaviors.

A zoospore is usually about 10 µm in size and has a kidney-like cell body [22, 23]. At least two unique traits distinguish zoospores from other prokaryotic and eukaryotic microswimmers currently studied using physical approaches: (i) The two flagella beat longitudinally along the anterior-posterior axis of the cell body and not laterally as in the case of the green algae *Chlamydomonas*; (ii) two flagella distinguished from each other as the anterior flagellum has a tinsel-like structure, while the posterior flagellum has a smooth whiplash one, both beating periodically with wave propagation directions outwards the body. These two flagella seem to be competing each other due to the opposite wave propagation directions. Contrarily, multiple mastigoneme structures on the anterior flagellum of zoospores were shown to have thrust reversal ability, which makes both flagella generate thrust in the same direction and propel the cell body forwards [19, 24]. Although it has been known about how zoospores swim, characteristics of the swimming and the beating flagella have not been statistically reported. The effects of mastigonemes on zoospore swimming also need to be carefully investigated since the mechanical properties of mastigonemes such as size, rigidity, density, can affect the swimming differently [25]. For instance, while mastigonemes are shown to generate thrust reversal in *P. palmivora* zoospores [24], they do not contribute to enhance swimming of *C. reinhardtii* [26].

In other microswimmers, their flagella are often synchronized to perform a cooperative swimming when they are a few microns away from each other [2, 27]. For examples, *C. reinhardtii* performs breaststroke swimming by two flagella drawing away and back to each other [4] or *E. coli, B. subtilis* bacteria’s flagella form a bundle and rotate together like a corkscrew to propel the cell body [3]. The question that whether zoospores, as eukaryotic swimmers, possess the similar cooperative behaviors of their flagella is of good interest. It was previously claimed that zoospore flagella are independent on each other and able to perform different tasks. Carlile [28] describes that the anterior flagellum is responsible for pulling the zoospore through water whereas the posterior flagellum acts as a rudder for steering the cell. However, Morris *et al*. [29] observe *P. palmivora* zoospores stop momentarily and then self-orientate their bodies to a new direction relatively to the posterior flagellum. Nevertheless, the cooperative actions of the motor and rudder were not carefully observed nor investigated, which remains unclear about how zoospores change direction either by random walks or in response to chemical and physical environment. This motivates us to unveil the physics behind individual swimming of zoospores.

In this article, we first investigate characteristics of zoospore trajectories at a global scale, then focus on the flagella scale’s swimming mechanisms. We observe that zoospores can perform long and stable straight runs, discontinued by active turning events. We obtain statistics of the trajectories and develop a numerical model to study and extrapolate the zoospore spreading characteristics solely by random walks. Then, we detail an in-depth study on the hydrodynamics of *P. parasitica*’s flagella and acquire a mathematical model to correlate the functions of two flagella on the motion of straight runs. Although theoretical models for microswimmers with single mastigonemes-attached flagella have been formulated [25, 30], models for microswimmers with two heterokont flagella have yet been considered as in case of zoospores. Here, we use Resistive Force Theory and further develop the model of a single flagellum with mastigonemes [25, 30] to adapt it with another smooth flagellum and a cell body, using a hypothesis of no interactions between two flagella. Moreover, we discover a unique active turning mechanism of zoospores including a body rotation then steering to a new direction, which results from the instantanous gait changing ability of their anterior flagellum. Our study reveals the mechanism and characteristics of zoospore spreading, which provides better insights on understanding and control of *Phytophthora* diseases.

## RESULTS AND DISCUSSION

### Characteristics of *P. parasitica’s* cell body and flagella

To understand the swimming, we first look at the cell body and flagellar structures of *P. parasitica*. By using Scanning Electron Microscopy (EM), we are able to observe the shape of the cell body and the positions of the flagellar base (Figure 1(B)). The cell in general has an ellipsoidal shape with tapered heads and a groove along the body. The size of the body is measured to be 8.8±0.4 µm (SEM) in length, and 4.7±0.1 µm (SEM) in width. The anterior flagellum attaches to the cell body in a narrow hole at one side of the groove, possessing an average length of 15.5 ± 0.1 µm (SEM). The posterior flagellum has the same diameter as the anterior’s (0.3 µm) but it is longer (20.3 ± 0.76 µm (SEM)), and attaches directly to the surface of the groove. The roots of two flagella are apart from each other with a distance of 2.9 ± 0.1 µm (SEM). Some mastigoneme structures were observed on the anterior flagellum, but could only be distinguished with difficulty by Scanning EM technique.

Observing the zoospores with Transmission Electron Microscopy (TEM) after negative staining, we discover multiple mastigonemes on both flagella (Figure 1(C1-3)). There are 2 different types of mastigonemes on the anterior flagellum: (type-1) straight and tubular shape, high density (*∼*13 per µm), 0.03 µm diameter, 1.5 µm long; (type-2) curved and irregular shape, longer and thicker in size (0.1 µm diameter, 1.8 µm long), and randomly distributed (Figure 1(C2)). The posterior flagellum instead has a smooth whip shape with plenty of very fine hairs on the surface (Figure 1(C3)). These hairs wrap around the flagellum to increase the contact surface, thus increasing the propulsion efficiency [31]. We also see a few type-2 mastigonemes on the posterior flagellum but they only appear near the root. The function of type-2 mastigonemes (Figure 1(C4)) is unknown, but their flexibility and random arrangement suggest that they might not contribute to generate drag. In contrary, the type-1 mastigonemes (Figure 1(C5)) are tripartite hairs that occur in most of Stramenopiles kingdom. They are known to be able to generate increased drag and reverse thrust for the anterior flagellum [24, 32].

### Statistics of individual swimming patterns

We investigate the characteristics of zoospore swimming by analyzing their trajectories and behaviors in water. To facilitate that, we perform microscopic assays where a low concentration of individual zoospores are released to an open thin film of water with thickness of *∼*100 µm on a glass slide. The setup of the water thin film can be visualized as a “swimming pool” that is not covered as we want to avoid the unwanted physical interactions of zoospores with the top when they experience aerotaxis. The zoospore swimming is captured at 60 fps (interval time between two consecutive frames, Δ*t ≈* 0.0167 s). The images are processed by Fiji [33] and Trackmate plugin [34] to semi-automatically track the positions of zoospores during the experiment duration (see Supp. Movie 1). Figure 2(A) illustrates the trajectories of zoospores captured from the microscopic assay. These trajectories indicate that zoospore can perform long and straight runs, some can even cross the whole observatory region. The straight runs are separated by multiple turning events when zoospores randomly change directions. With this swimming strategy, zoospores can be categorized as run-and-tumble active particles [35]. From the position data of each zoospore over time, we achieve its movement characteristics defined by two parameters: magnitude of speed *U* and moving directions *θ* (Figure 2(B-C)), after applying moving average method with step length *n* = 12 to improve the accuracy of moving direction and instantaneous speed of the zoospore. From *U* values, we can separate the movement of zoospores into 2 states: running state during straight runs and stopping state at turning events. While running, *U* and *θ* vary around a constant value. At turning events, *U* drops drastically, (occasionally close to 0) then quickly recovers, *θ* also rapidly changes to a new value. *U*_th_ is defined as the threshold speed that separates the two states of running and turning. With *U*_th_, we determine two important parameters of zoospore swimming: (1) running time *τ*_r_ as the duration when *U ≥ U*_th_, deciding how long a zoospore is able to travel without turning; (2) stopping time *τ*_s_ as the duration, when *U ≤ U*_th_, for a zoospore to perform a turn.

**FIG. 2.**
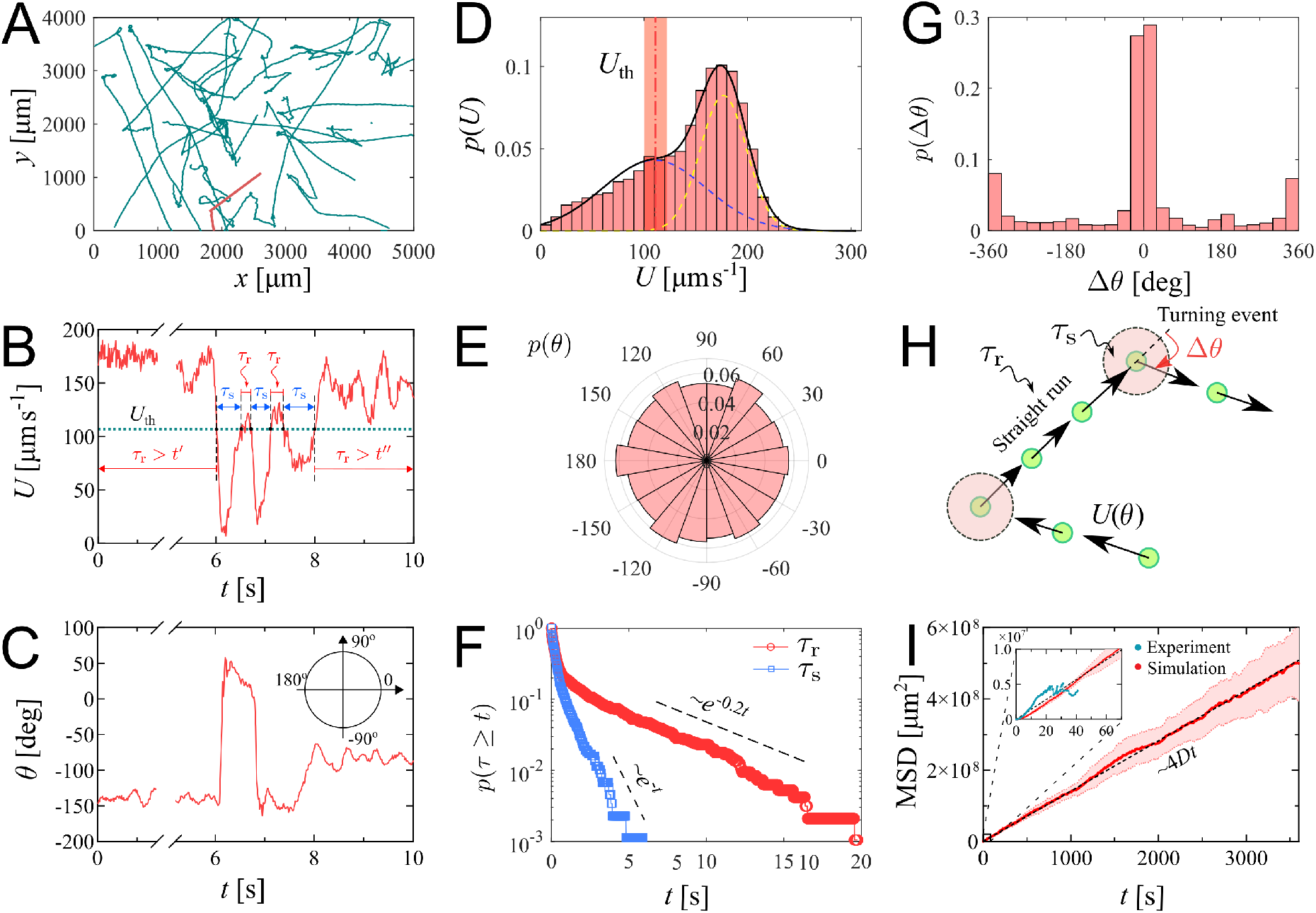
Swimming trajectories of *P. parasitica* zoospores. (A) Trajectories of zoospores swimming in water captured from the microscopic assay for 60 s. Sample size *N* = 58. Note: not all trajectories are shown. Each position of the zoospores is captured every Δ*t* = 0.0167 s. The trajectories are smoothed with moving average (step length *n* = 12). (B) The progression of speed *U* and (C) moving directions *θ* over time of a single zoospore extracted from the population in the assay. (D) Distribution of zoospore speed *p*(*U*). (E) Polarity distribution of moving direction *p*(*θ*). (F) Survival curves *p*(*τ ≥ t*) of the running time *τ*_r_ and stopping time *τ*_s_. (G) Distribution of turning angle *p*(Δ*θ*). (H) Schematics showing the strategy of the simulation model of zoospores swimming in water. (I) The estimated mean squared displacement (MSD) over time intervals *t*, constructed from the simulation data. The inset compares the experimental data and simulation of MSD at the experimental time-scale of 60s. By simulation, at long time scale of 1 hour, MSD of zoospores shows a diffusion of Brownian particles with the diffusion coefficient *D* = 3.5 × 10^−4^cm^2^ s^−1^.

We plot the distribution of *U* for all the trajectories of zoospores swimming in the observatory region for duration of 60 s (total number of zoospores *N* = 58) in Figure 2(D). The speed distribution *p*(*U*) exhibits a combination of two different normal distributions: 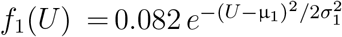 with µ = 176.6, *σ*_1_ = 22.3 µm s^−1^, and 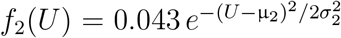 with µ_2_ = 110, *σ*_2_ = 53 µm s^−1^. This bimodal distribution of *U* indicates that zoospore speed fluctuates around two speed values *U* = µ_1_ and *U* = µ_2_, corresponding to two behavioral states of running and turning. The distribution *f*_1_ is associated with the running state where zoospores experience stable moving speed, while *f*_2_ represents the speed at turning events where zoospores reduce their speed from a stable running speed to 0 then quickly recover. We achieve the fitting curve for *p*(*U*), resulting from the sum of two Gaussian fits *f*_1_ + *f*_2_, and choose the speed at inflection point of the fitting curve where µ_s_ *≤ U ≤* µ_r_ as the speed threshold to separate two behavioral states, *U*_th_ = 111.5 µm s^−1^. *U*_th_ at the inflection point determines a turning point for a significant change of the speed from running to turning state. The sensitivity of our *U*_th_ selection can be tolerated by ±10 % of the chosen value, ranging from 100 to 122.5 µm s^−1^ (See Supp. Materials). We also plot the polarity distribution of zoospores in Figure 2(E) based on the moving direction *θ* and acquire an equally distributed in all directions. With the defined *U*_th_, we calculate and plot the distributions of *τ*_r_ and *τ*_s_ in form of survival curves *p*(*τ ≥ t*) in Figure 2(F). Both survival curves show complex behaviors of zoospores during running and turning. The statistics *p*(*τ*_s_ *≥ t*) can be considered as sum of two exponential decays with average stopping time 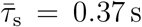. Also, *p*(*τ*_r_ *≥ t*) is in form of two exponential decays with average running time *τ*__r_ = 1.0 s. We can estimate that zoospores stop and turn with the frequency 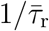 greater than 1 Hz (*p*(*τ*_r_ *<* 1.0 s) = 0.82). Based on the moving direction over time, we calculate the average turning speed of zoospores at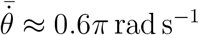. At each stopping time they perform a turning angle Δ*θ* = *θ*_i_ *−θ*_e_, where *θ*_i_ and *θ*_e_ is the moving direction right before and after each turning event, respectively. The distribution of Δ*θ* is shown in Figure 2(G), demonstrating the equal preference of turning directions (with positive angle values indicating counter-clockwise, and negative as clockwise direction). It is also shown that zoospores preferentially turn with the angle around 0^*°*^, which we speculate that it results from the failed out-of-plane movement when zoospores swim near the water/air interface during their aerotaxis (See Supp. Movie 2).

Since the motion of zoospore is characterized by the succession of straight runs and turning events, as illustrated in Figure 2(H), in order to quantify their large-scale transport properties, we assemble all previous measurements in the following way. Each straight run is characterized by a speed *U*_r_ *> U*_th_ and a duration *τ*_r_ drawn from the distributions in Figure 2(D) and Figure 2(F), respectively. After a run phase, an idle phase of duration *τ*_s_, drawn from Figure 2(F) follows. The moving direction of the *r*-th run phase is given by 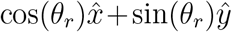, i.e. it is parameterized by an angle *θ*_*r*_. Note that moving direction of two consecutive *r*-th and (*r*+1)-th run phases are correlated. Moreover, *θ*_*r*+1_ = *θ*_*r*_+Δ*θ*, where Δ*θ* is a random angle drawn from the distribution in Figure 2(G). Mathematically, the position **x**_*m*_ of the zoospore after *m* run phases is given by 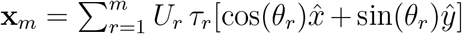, and its mean-square displacement is MSD(*m*) =⟨(**x**_*m*_*−(***x**_*m*_))^2^⟩, where ⟨…⟩ denotes average over realizations of the process. Our simulation results in a diffusive behavior of zoospores, with the MSD proportional to *t* (see Supp. Movie 3). The diffusion coefficient is then obtained from 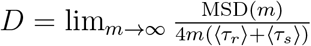. In the computation of MSD and *D*, we assume that the only random variable exhibiting correlations is *θ*_*r*_, while *U*_*r*_, *τ*_*r*_, and *τ*_*s*_ are uncorrelated. This procedure allows us to obtain a reliable estimate of *D*, with *D* = 3.5 × 10^−4^cm^2^ s^−1^, which is in the same order of magnitude as the diffusion coefficient of *C. reinhardtii* ‘s [4]. We stress that direct measurements of *D* based on experimental MSD data are highly unreliable given the relatively small number of trajectories and their short duration. In our case, the data are enough to obtain reliable estimates on the distributions of Δ*θ, U*_*r*_, *τ*_*r*_, and *τ*_*s*_ from which, as explained above, *D* can be reliably estimated from simulations. Similar methods of using a theoretical random walk model to estimate the macroscopic parameters from the microscopic experiment have been previously developed for *C. reinhardtii* [36, 37]. We emphasize that *D* represents the estimation of diffusion coefficient of individual swimming of zoospores from random walk process. This is more to show the intrinsic ability of individual zoospores to perform spatial exploration, rather than to quantify the bulk diffusivity where the collective swimming behaviors, which involve zoospore-zoospore interactions, play a major role.

### Role of two flagella in swimming motions of zoospores

Our statistics study on swimming trajectories of zoospores has delivered characteristics of their movement at large scale, including the straight runs and turning events. These motions are controlled by two flagella oriented in opposite directions along the cell body’s anterior-posterior axis. In this section, we look in-depth to how these two flagella together generate speed and perform turning for zoospores by conducting microscopy assays at small length scale, of which the flagella are visible and in very short time scale.

#### Straight runs

We record movement of zoospores during their straight runs with visible flagella by conducting brightfield microscopy with 40× objective and a high-speed camera capturing at 2000 fps at exposure time 200 µs. In Figure 3(A), we show images of a *P. parasitica’s* zoospore swimming by two flagella beating in sinusoidal shapes with the wave propagation in opposite directions. While translating, the cell body gyrates around the moving direction simultaneously, which results in a helical swimming trajectory (see Supp. Movie 4 for a long run of a zoospore swimming in water). We believe that this gyrational motion might result from the intrinsic chiral shape of the zoospore body and off-axis arrangement of their flagella (Figure 1(B)). Indeed, previous studies have shown that chirality of a microswimmer’s body induces spontaneous axial rotation resulting from the translational motion [38–40]. From multiple observations, we obtained the pitch and radius of the helical trajectories at *p* = 130 ± 8 µm (SEM) and *R* = 4.0 ± 0.2 µm (SEM), respectively (data presented in Supp. Materials). We then estimate the gyrational speed of the cell body 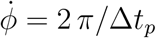, where Δ*t*_*p*_ is the duration the zoospore travels through a full turn of the helical path (Figure 3(B)). We obtain 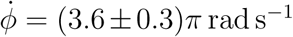 (SEM) (Supp. Materials). The observations of helical trajectories also confirm that each flagellum of zoospores beat as a flexible oar in a 2D plane as we observe the two flagella flattened into two straight lines during the gyration. Thus, zoospores are not expected to swim in circles when interacting with no-slip boundaries as seen in case of *E. coli* with a rotating flagellar bundle. The “curved straight runs” of zoospores that we observed in Figure 2(A) might result from rotational diffusion and thermal fluctuations.

**FIG. 3.**
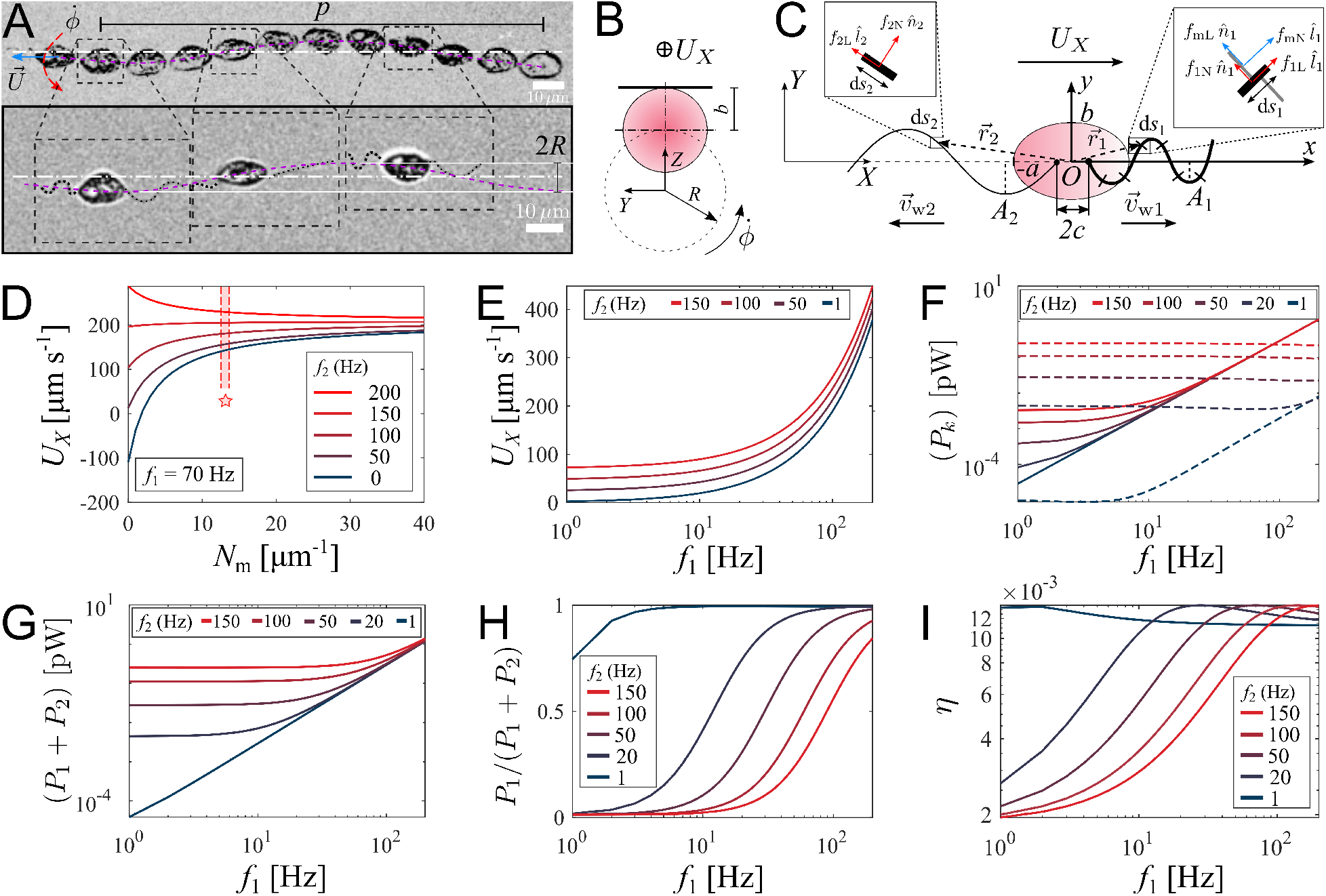
Theoretical model of swimming individual *P. parasitica* zoospore. (A) Images of an individual zoospore swimming with two flagella beating in sinuisoidal waveform shapes and its cell body gyrating with rate 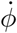 while moving forward with speed *U*. The combined motion results in a helical swimming trajectory with pitch *p* and radius *R*. (B) Schematics showing the gyration of the cell body. (C) Theoretical model of a zoospore translating in a 2D plane using Resistive Force Theory. (D) The dependence of translational speed *U*_*X*_ on the type-1 mastigoneme density (*N*_m_). The range of *N*_m_ with symbol (*\**) indicates the values measured by TEM. (E) The effects of beating frequencies of the two flagella, *f*_1_ and *f*_2_, on zoospore speed *U*_*X*_, (F) power consumption of each flagellum *P*_*k*_, (G) total power consumption of two flagella (*P*_1_ + *P*_2_), (H) power distributed to the anterior flagellum and (I) propelling efficiency of both flagella *η*. In these plots, *N*_m_ is set at 13 µm^−1^.

We retrieve the parameters of the beating flagella including beating frequencies *f* and wavelengths *λ*, by applying kymographs on the cross-sections within the two flagella and normal to the moving direction of the zoospore. We present more details of the parameter retrieval with kymographs in Supp. Materials. Our data show that when swimming in straight runs, the anterior flagellum of zoospores usually beats at *f*_1_ *≈* 70 Hz while the posterior flagellum beats with the frequency *f*_2_ *≈* 120 Hz that is approximately 1.7-fold faster than the anterior’s. We focus on a very short duration of less than 50 ms, which is equivalent to a translation of less than 10 µm. Compared with the rotation motion at time-scale of 1 s, we neglect the effect of the cell body’s rotation in this short duration. We then develop a mathematical model to study how the dynamics of beating flagella helps generating thrust for the cell to move forward. We assume that the gyration of the cell body does not affect the shapes and motions of the flagella since the beating frequencies of the two flagella are much higher than the gyrational speed. The gyration also does not contribute to the translation as we consider it as a passive motion resulting from the chirality of zoospores. Thus, the swimming zoospore can be considered as a 2D model (Figure 3(C)) in which the cell body is an ellipse defined as *x*^2^*/a*^2^ + *y*^2^*/b*^2^ = 1 in its body-fixed frame (*xOy*), with the anterior and posterior flagellum having sine waveform shapes defined as

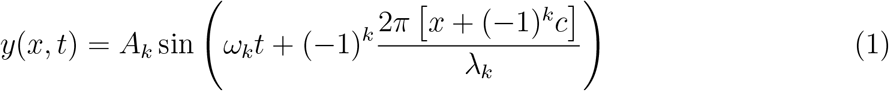

as (−1)^*k*^*x ≤ −c*, where *A*_*k*_ is the amplitude, *ω*_*k*_ is the angular speed, *λ*_*k*_ is the wavelength of the anterior (*k* = 1) and the posterior flagellum (*k* = 2). The two flagella are attached to the cell body at two points both lying on *x*-axis and distanced to the origin *O* a gap of *c*, and beating with the wave propagation *-v*_w_ having directions against each other. The anterior has length *L*_1_ and diameter *d*_1_ while those of the posterior are *L*_2_ and *d*_2_. Additionally, there are multiple tubular type-1 mastigonemes with length *h* and diameter *d*_m_ attached to the surface of the anterior flagellum with density *N*_m_ indicating the number of mastigonemes attached on a unit length of the flagellum. It is important to determine the flexibility of these mastigonemes as it would impact the ability of the mastigonemes to produce drag. We estimate the flexibility by a dimensionless parameter, which were carefully characterized in previous studies [25, 41], 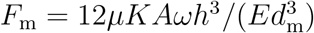, where *µ* is the fluid viscosity, *K* = 2*π/λ* is the wave number, *E* is the Young modulus of the mastigonemes. With this estimation, if *F*_m_ *<* 0.1, mastigonemes are considered as fully rigid. In case of zoospores’ mastigonemes, we achieve *F*_m_ at order of 10^−4^, which is much lower than 0.1. Thus, we can assume that the mastigonemes of zoospores are non-deformable and rigidly attached to the anterior flagellum. As a result, hydrodynamic interactions between neighboring mastigonemes can also be neglected. Additionally, we ignore the effects of the type-2 mastigonemes in producing drag due to their non-tubular and random structures.

Zoospores swim in water with very low Reynolds number (*Re <<* 1), resulting in negligible inertia, dominant viscous force and the kinetic reversibility [42, 43]. Microswimmers with flexible flagella generate thrust from drag force acted by fluid on the flagellum segments. In our model, we use Resistive Force Theory (RFT) to deal with the calculation of fluid’s drag force on the two flagella of the zoospore. RFT has proven to be an effective and accurate method to predict the propulsive force and velocity of microswimmers regardless of the interactions of flagellum-flagellum or flagellum-body [44–46]. In case of zoospores where two flagella are in opposite directions, and the flagellum-body interaction is insignificant, RFT is a suitable solution to apply. Following this method, each flagellum is divided into an infinite numbers of very small segments with length d*s*_*k*_, and each segment is located in the body-fixed frame (*xOy*) by a position vector *-*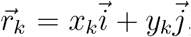, where 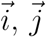 are the unit vectors in *x*- and *y*-direction, respectively; *x*_*k*_ and *y*_*k*_ satisfy the shape equation (Equation 1) for the anterior (*k* = 1) and the posterior (*k* = 2).

RFT states that the drag force by fluid acting on an infinitesimal segment d*s* of the flagellum is proportional to the relative velocity of fluid to the flagellum segment [44, 47, 48], as follows

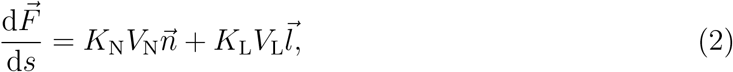

where *V*_N_ and *V*_L_ are two components of relative velocity of fluid in normal and tangent direction to the flagellum segment, *K*_N_ and *K*_L_ are the drag coefficients of the flagellum in normal and tangent to the flagellum segment, 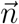 and 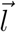 are the unit vectors normal and tangent to the flagellum segment, respectively. The drag coefficients *K*_*N*_ and *K*_*L*_ are estimated by Brennen and Winet [49], which depends on fluid viscosity, the wavelength and diameter of a flagellum.

We then apply RFT on each flagellum of the zoospore to calculate the total drag force acting on it. For the posterior flagellum, each segment d*s*_2_ is a simple smooth and slender filament (see inset d*s*_2_ in Figure 3(C)), having drag coefficients *K*_N2_ and *K*_L2_. For the anterior flagellum, each segment d*s*_1_ contains additional *N*_m_d*s*_1_ mastigonemes. Using a strategy from previous models for flagella with mastigonemes [25, 30], we consider these mastigonemes stay perpendicular to the segment itself (see inset d*s*_1_ of Figure 3(C)) and also act as slender filaments experienced drag from water. Interestingly, due to the direction arrangement, the relative velocity normal to the flagellum segment results in drag force in tangent direction to the mastigonemes, and subsequently, the relative velocity tangent to the flagellum segment results in drag force in normal direction to the mastigonemes. In another perspective, we can consider the anterior flagellum receives additional drag from the mastigonemes, which is presented by two increased drag coefficients in normal and tangent direction defined as

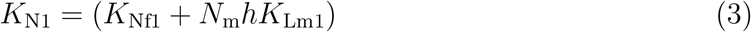

and

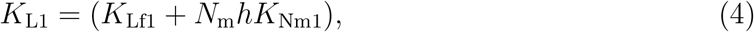

respectively. Here, *K*_Nf1_ and *K*_Lf1_ are the drag coefficients in normal and tangent direction of the flagellum filament, respectively; *K*_Nm1_ and *K*_Lm1_ are the drag coefficients in normal and tangent direction of the mastigonemes, respectively.

In low Reynolds number condition, total forces equate to zero due to approximately zero inertia. Thus, we derive translational velocity *U*_*X*_ of the zoospore as shown in Equation 5

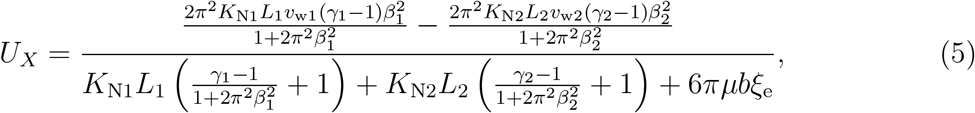

where *v*_wk_ = *λ*_*k*_*f*_*k*_ is the wave propagation velocity of the flagellum, *γ*_*k*_ = *K*_Lk_*/K*_Nk_ is the drag coefficient ratio, *β*_*k*_ = *A*_*k*_*/λ*_*k*_ is the flagellar shape coefficient, and *ξ*_e_ is the shape coefficient of the ellipse cell body. See Supp. Materials for the detailed derivatives.

We first study the effects of mastigonemes on zoospore speed by ploting the value of *U*_*X*_ with different density of the mastigonemes, and varying the beating frequency of the posterior flagellum *f*_2_ while the anterior flagellum beats with a usual *f*_1_ = 70 Hz in Figure 3(D). The dimensions and physical parameters of the zoospore’s cell body and two flagella are taken from Supp. Materials. The plot shows that the appearance of mastigonemes results in a reversed thrust from the anterior flagellum. To illustrate this, when there are no mastigonemes on the front flagellum (*N*_m_ = 0) and the posterior flagellum is excluded (*f*_2_ = 0, *L*_2_ = 0), the anterior flagellum generates a thrust in negative *X*-direction (*U*_*X*_ *<* 0). This agrees well with previous studies modeling a smooth reciprocal beating flagellum [44, 45, 47]. But since *N*_m_ *>* 2, the velocity of the zoospore is reversed to positive *X*-direction due to the extra drag from the mastigonemes. This phenomenon was also described in hydrodynamics by Namdeo *et al*. [25], and observed in experiment of Cahill *et al*. [24]. Interestingly, when changing directions, zoospores are observed to swim with the solely beating anterior flagellum to pull the body forwards while the posterior flagellum is immobile (Figure 4(A)) and Supp. Movie 5), which confirms the thrust reversal ability of the mastigonemes. The beating frequency of the anterior flagellum in this case increases to *f*_1_ *≈* 110 Hz. This finding also reassures the importance of mastigonemes on zoospore swimming, which is not similar to those of *C. reinhardtii* [26]. The front flagellum of zoospores has fibrillar mastigonemes, similarly to *Chlamydomonas*, but at higher density with tubular shape and larger in size, that could render into account of the different beating properties from the smooth posterior flagellum. However, high mastigoneme density (more than 20 per 1 µm flagellum length) shows mild effect on speed. From TEM images taken at the anterior flagellum, we estimate the mastigoneme density by averaging the number of mastigonemes manually counted over a flagellum length (See Supp. Materials). We obtain *N*_m_ = 13.0 ± 0.8 µm^−1^ (SD), which falls between the optimum range to generate speed.

**FIG. 4.**
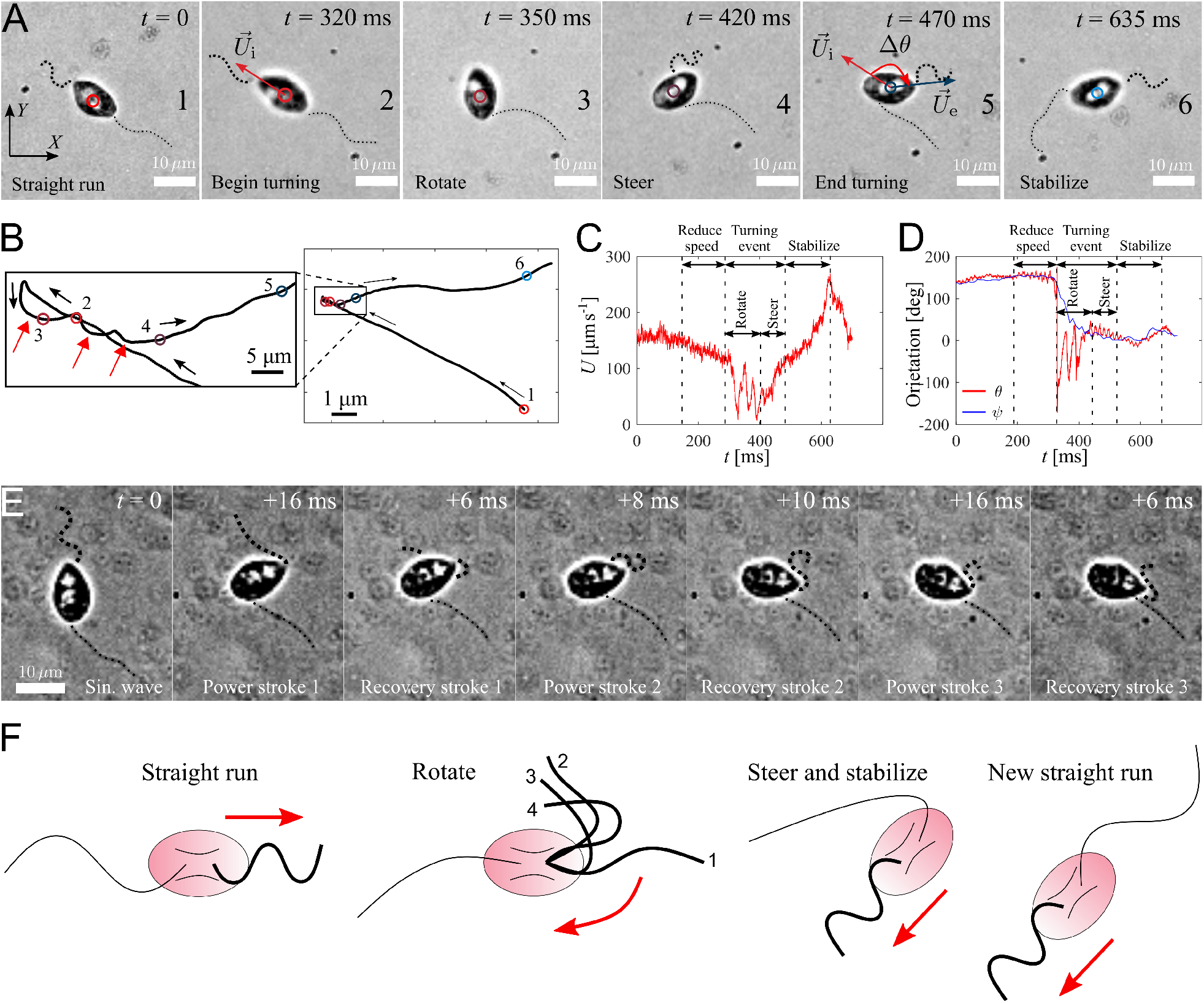
Active turning of individual *P. parasitica* in water. (A) Images of a zoospore changing direction. The two flagella cooperate to help the cell body rotate and steer to a new direction achieving a turning angle Δ*θ*. (B) Trajectory of the zoospore during the turning event. Three red arrows represent 3 back and forth stroke-like motions. (C) The speed *U* of the zoospore during the turning event. The turning starts when the speed begins to fluctuate with large magnitude and lower frequency, and lasts for a duration of *τ*_s_ with a rotation of the cell body followed by steering to the new direction. (D) The moving directions *θ* and the body orientation *ψ* of the zoospore during the turning event. (E) Images of the anterior flagellum of a zoospore beats with power and recovery stroke, similarly to *C. reinhardtii* ‘s in a temporal zoom corresponding to the “Rotate” step of the turning event (but not from the same movie as Figure 4(A)). (F) Schematics to describe the gait of the flagella during a turning event. (1-2) Power stroke 1, (2-3) recovery stroke 1, (3-4) power stroke 2.

To understand how the coordination of two flagella influences zoospore speed, we vary the beating frequency of one flagellum while the other’s remains constant and obtain the resultant speed (Figure 3(E)). We find that although both flagella contribute to zoospore speed, the anterior flagellum has larger impact on speed than the posterior one. For instance, the anterior flagellum can singly generate a speed 3-fold higher than the posterior flagellum can do at the same beating frequency. Moreover, the additional speed contributed by the anterior flagellum remains almost the same regardless of the varation of frequency of the posterior flagellum, while the speed contribution of the posterior flagellum decreases as the anterior flagellum increases its frequency. Since the contribution to zoospore speed is different between two flagella, we ask whether the energy consumption of each flagellum might also be different or equally distributed. In our model, each flagellar segment consumes a power deriving from the dot product between the drag force of water acting on the segment and the relative velocity of water to the segment, which can be written as 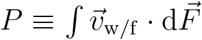 (see Supp. Materials for detail derivatives). We achieve that, the power consumption of each flagellum depends on both flagella at low frequencies (*f ≾* 40 Hz), and becomes solely dependent on its own beating at high frequencies (*f ≿*40 Hz) (Figure 3(F)). At high frequencies, the anterior flagellum consumes approximately 5-fold more power than the posterior, given the same beating frequency. As a result, the total power consumed for both flagella becomes less dependent on the posterior flagellum as the anterior flagellum beats faster (Figure 3(G)). In Figure 3(H), we show the fraction of power consumed by the anterior flagellum over the total power consumption of the zoospore. We notice that, at the same beating frequency with the posterior flagellum, the anterior flagellum accounts for *∼*80 % of the total power. Interestingly, when the frequency of the posterior flagellum is *∼*1.7-fold higher than that of the anterior flagellum, both flagella consume the same amount of power. Indeed, this result agrees well with our experimental data, in which we obtain that the anterior flagellum normally beats at 70 Hz and the posterior flagellum at 120 Hz. Thus, we can speculate that the energy is equally distributed for both flagella. In addition, we also estimate the propelling efficiency of the zoospore 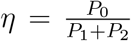, with *P*_0_ as required power to move the cell body forward at speed *U*_*X*_ (see Supp. Materials for derivatives). We achieve that *η* is higher as the anterior flagellum increases its frequency (Figure 3(I)). The efficiency reaches its maximum value at *∼*1.2 % when the beating frequency of the posterior is *∼*1.7-fold higher than that of the anterior flagellum. Overall, we show that the energy is shared in comparable manner between two flagella, but the anterior flagellum has more influence on zoospore speed, power consumption and propelling efficiency. On the other hand, the posterior flagellum provides a modest contribution to zoospore speed despite beating at higher frequency and consuming half of the energy. Taking the fact that the anterior flagellum can beat singly with the posterior flagellum being immobile during turning events, we can speculate that the anterior flagellum is the main motor of zoospores. Nevertheless, the function of the posterior flagellum remains cryptic and we investigate more on its role during turning events.

#### Turning events

We capture zoospores changing their directions by a unique active turning mechanism and an interesting coordination between two flagella. The movies of zoospore turning are recorded at 2000 fps with the same setup as the microscopic assay (Figure 4(A) and Supp. Movie 5). Using Fiji and Trackmate, we track the positions of the zoospore and plot the trajectory, smoothed by moving average with step length *n* = 40, during its turning event (Figure 4(B)). We then compute the zoospore speed *U* (Figure 4(C)), and the moving directions *θ* from the trajectory while manually measure body orientation *ψ* over time (Figure 4(D)). We observe that at first, to prepare for the turn, the zoospore reduces its speed as both flagella beat with smaller amplitudes. Then, the posterior flagellum instantaneously stops beating, which marks the beginning of a turning event (Figure 4(A1-2)). The anterior flagellum now takes full control of zoospore motion during turning. The zoospore then perform two distinct sequences of motions right after the posterior stops beating: (i) rotation of the cell body out of the old direction resulting from a few repetitive stroke-like beating of the anterior flagellum (Figure 4(A3)), then (ii) steering towards a new direction as the anterior flagellum switches back to normal beating with sinusoidal waveform to propel the body (Figure 4(A4-5)). These two sequences of turning can be distinguished by two different patterns of trajectory during turning. While the rotation is indicated by multiple circular curves, in which each of them corresponds to a stroke-like back and forth beating motions, the steering results in straight line (Figure 4(B)). Additionally, the zoospore speed *U* during this period also consists of two different patterns of fluctuations (Figure 4(C)), while the moving direction *θ* changes from large fluctuations (for rotation) to stable direction (for steering) (Figure 4(C)). The turning event ends when the cell body stably moves in the new direction, which is indicated by the recovery of speed and the overlap of the moving direction and body orientation. The anterior flagellum continues to propel the cell body out of the location of the turning event (*∼*5 µm away), which we call ”stabilize“ step (Figure 4(A6)). We notice that in this step, the anterior flagellum beats at a higher frequency than usual (*∼*110 Hz, compared to normally at *∼*70 Hz) and achieves a high-speed of *∼*250 µm s^−1^. Finally, the posterior flagellum resumes its beating and the zoospore returns to the normal straight run state.

It is striking that the anterior flagellum is able to completely change its gait from sinu-soidal traveling wave to stroke-like beating during the rotation step of the turning event. Thus, we need to have experimental evidence from direct observation of the shape of the anterior flagellum during this gait changing period, which is sometimes hidden underneath the cell body due to the off-axis postion of the flagellar base, to extend our understanding on this turning behavior. We capture new movies of zoospore turning events, but the focus plane is slightly offset from the focus plane of the cell body (Figure 4(E) and Supp. Movie 6). We observe that the anterior flagellum instantaneously perform continuous power and recovery strokes, similarly to the breast-stroke beating of *Chlamydomonas*’s flagella. As a result, the cell body can rotate quickly about a fixed point, which helps the zoospore make a sharp turn. We summarize the turning of zoospores by a schematic in Figure 4(F). Indeed, this rotation motion is similar that of uniflagellate *C. reinhardtii* [50], which strengthens the evidence for the actively switchable beating pattern of zoospores’ anterior flagellum. While the anterior flagellum plays a major role in turning events on top of straight runs, the posterior flagellum only stays immobile throughout the process. Here, we can confirm that the posterior flagellum does not act as a rudder to steer the direction. Instead, it is fully stretched during turning events and might contribute to increase drag at one end of the cell body. The function of the posterior flagellum is not completely clear regarding thrust production during translation or turning. Thus, we speculate that it might contribute to other non-physical activities such as chemical and electrical sensing. The advantage of zoospore turning mechanism is that it allows them to actively and quickly achieve a new direction, which is not the case of other microswimmers such as the tumbling of *E. coli* ‘s [51, 52] or *Chlamydomonas*’s [4, 53].

## CONCLUSION

We have performed the first systematic study of the swimming pattern and spreading features of *P. parasitica* zoospores, a plant pathogen, which is considered a major agricultural threat. Combining high-speed imaging and Resistive Force Theory, we show how the two opposite flagella are coordinated in producing thrust by beating together, allowing the microorganism to achieve high-speed swimming during straight runs. Furthermore, we find that turning is a coordinated process, in which the the posterior flagellum stops beating, while the anterior flagellum actively moves causing the cell body to rotate. Finally, we explain how fast-swimming periods and active turning events combine to produce a diffusion coefficient of *D* = 3.5×10^−4^ cm^2^ s^−1^, a quantity that characterizes spatio-temporal spreading of this pathogen during plant epidemics.

It is worth stressing the motility pattern exhibited by the zoospores represents an Eukaryotic version of the “run-and-tumble” motility class exhibited by bacteria peritrichous bacteria. Several eukaryotic swimmers, e.g. spermatozoa, do not exhibit such kind of motility pattern, but *C. reinhardtii* have been reported to also fall into this category [4]. There are, however, important differences. *Chlamydomonas* possess two identical flagella located at one tip of the cell. Reorientation events occur during asynchronous beating periods of the two identical flagella, while straight runs require synchronous beating. In sharp contrast to this picture, we show that in zoospores while straight runs also involve a coordinated beating of the opposite and different flagella, turning involves temporary halting of posterior flagellum, while the anterior flagellum continues beating. This strongly suggests that *P. parasitica* navigates using a fundamentally different internal regulation mechanism to control swimming, than *C. reinhardtii*, a mechanism that is likely to be present in other Eukaryotic swimmers with two opposite and different flagella.

We believe our findings on the coordination of two flagella bring more insights on zoospore swimming dynamics. It was also not known from the literature about the role of each flagellum on the straight runs, and on turning events in particular. We show that although the energy is shared in comparable manner between both flagella, the anterior flagellum contributes more to zoospore speed. Zoospores actively change directions thanks to the sole beating of the anterior flagellum. We believe that anterior flagellum could be the main motor of zoospores that is in charge of generating speed, changing beating patterns from sinusoidal wave to power and recover stroke to quickly rotate the body, while the posterior flagellum might play a role in chemical/electrical sensing and providing an anchor-like turning point for zoospores, instead of acting like a rudder as previously hypothesized. These findings pave new ways for controlling the disease since now we can have different strategies on targeting one of the flagella.

## EXPERIMENTAL SECTION

### *P. parasitica* mycelium culture and zoospore release

We culture mycelium of *Phytophthora parasitica* (isolate 310, *Phytophthora* INRA collection, Sophia-Antipolis, France) [18, 19] routinely on malt agar at 24^*°*^C in the dark. To produce zoospores, we prepare the mycelium which is grown for one week in V8 liquid medium at 24^*°*^C under continuous light. The material was then drained, macerated and incubated for a further four days on water agar (2%) to induce sporangiogenesis. Zoospores are released from sporangia by a heat shock procedure. We place a petri-dish of mycelium inside a refrigerator at 4 ^*°*^C for 30 minutes, then pour 10 ml of 37 ^*°*^C distilled water on top of the mycelium and continue to incubate it at room temperature (25 ^*°*^C) for another 30 minutes. Zoospores escape from sporangia and swim up to the water. The zoospore suspension is then collected for further experiments.

### Scanning Electron Microscopy and Transmission Electron Microscopy with negative staining

For Scanning Electron Microscopy and Transmission Electron Microscopy, cell pellets are fixed with in a 2.5 % glutaraldehyde solution in 0.1 M sodium cacodylate buffer (pH 7.4) at room temperature (*∼* 25 ^*°*^C) for 1 hour and then stored at 4 ^*°*^C. For Scanning EM observations, after three rinsing in distilled water, protists are filtered on a 0.2 µm isopore filter. Samples on filters are subsequently dehydrated in a series of ethanol baths (70 %, 96 %, 100 % three times, 15 minutes each). After a final bath in hexamethyldisilazane (HMDS, 5 minutes), samples are left to dried overnight. Samples on filters are mounted on Scanning EM stubs with silver paint and coated with platinum (3 nm) prior to observing. The Scanning EM observations are performed with a Jeol JSM-6700F scanning electron microscope at an accelerating voltage of 3 kV.

For TEM observations, samples are prepared using the negative staining method. After three rinsing in distilled water, a drop of cells suspension (*∼*10 µl) is left for 5 minutes on a TEM copper grid (400 mesh) with a carbon support film. The excess liquid is removed with a filter paper. Subsequently, staining is done by adding a drop of 0.5 % (w/v) aqueous solution of uranyl acetate on the grid for 1.5 minute, followed by removal of excess solution. The TEM observations are carried out with a JEOL JEM-1400 transmission electron microscope equipped with a Morada camera at 100 kV.

### Microscopic assays of zoospores

We pipette a droplet of 10 µl water containing zoospores onto a microscopic glass slide and spread the droplet to thoroughly cover the marked area of 1 × 1 cm and become a thin film of approximately 100 µm thickness. We do not cover the droplet to prevent the unwanted interactions between the zoospores and rigid surface of coverslips. We observe the swimming of individual zoospores inside the flattened droplet under a bright field transmission microscope (Nikon Eclipse T*i* 2, Minato, Tokyo, Japan) at 40× objective with the high speed camera Phantom v711 (Vision Research, NJ, USA). For the experiment to observe the swimming trajectories of the zoospores, we use 4× objective to capture a large swimming region of 5000 × 4000 µm. The captured images are processed by Fiji with the Trackmate plugin.

### Estimation of trajectory parameters

An individual zoospore positions are captured at each time frame Δ*t*. At each *t*_*j*_ = *j*Δ*t* (*j* = 1, 2, 3, …), the zoospore has a position *z*_*j*_ = (*x*(*t*_*j*_), *y*(*t*_*j*_)). First, we smooth the trajectory by moving average with step *n*. The smoothed positions *Z*_*j,n*_ are calculated as

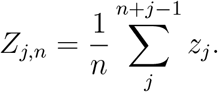

Each new position *Z*_*j,n*_ possesses a velocity vector with speed

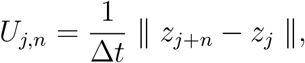

and moving direction (angle between velocity vector and *x*-axis)

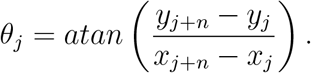

## ACKNOWLEDGMENTS

This work has been supported by the French government through the UCA^JEDI^ Investments in the Future project managed by the National Research Agency with the reference number ANR-15-IDEX-01. CCMA electron microscopy equipments have been funded by the Région Sud - Provence-Alpes-Côte d’Azur, the Conseil Général des Alpes Maritimes, and the GIS-IBiSA. We thank Xinhui Shen for fruitful discussion, as well as Marcos, David Gonzalez-Rodriguez for their careful reading and giving valuable feedback to improve the manuscript.

## Notes

### Competing Interest Statement

The authors have declared no competing interest.

### Summary of Updates

Writing improvement for section "Straight runs", "Turning events" and "Conclusion". Figure 3F-I added. Figure 4E-F added. 2 new supp. movies added.

