## Supplementary Materials for "Coordination of two opposite flagella allows high-speed swimming and active turning of individual zoospores"

#### 1 Obtain characteristics of beating flagella by kymograph

We first use Fiji with StackReg plugin to fix the position of the cell body in the swimming movies, and only the two flagella beating.

We retrieve the parameters of the beating flagella by applying kymographs on multiple cross-sections which are along to or normal to the moving direction at the positions shown in Fig. S1. These kymographs help converting the complex time-dependent 2D image data of the flagella from $(X, Y, t)$ to separate signals $(X, t)$ and $(Y, t)$. As a result, the kymographs at cross-section (1) and (2) give us the average amplitudes $A$ of the waveform shapes of the two flagella, and the ones at (3) and (4) provide us with information of beating frequencies $f$ and wavelengths $\lambda$.

#### 2 Variation of $U_{\mathrm{th}}$

To verify if the way we chose the value of $U_{\mathrm{th}}$ will affect the result of turning angles and other swimming parameters, we vary the $U_{\mathrm{th}}$ value by $\pm 10\,\%$ of the chosen value, which ranges from 100 to 122.5 $\mu\mathrm{m\,s^{-1}}$. We plot the running time $\tau_{\mathrm{r}}$, stopping time $\tau_{\mathrm{s}}$ and turning angles $\Delta\theta$ according to the varied $U_{\mathrm{th}}$. Data are shown in Figure S2. We see that the changes do not affect the results of $\tau_{\mathrm{r}}$, $\tau_{\mathrm{s}}$ and $\Delta\theta$ estimation. Thus, we believe the sensitivity of $U_{\mathrm{th}}$ selection can be tolerated by $\pm 10\,\%$.

#### 3 Characteristics of helical trajectories

From the microscopic assays with high-speed camera and 40× objective, we observe zoospores swimming in helical trajectories. We manually measure the characteristics of these helical trajectories and present the data in the table below.

*Present address: Institut Jacques Monod (IJM), University of Paris, CNRS, UMR 7592, 75013 Paris, France
†
‡

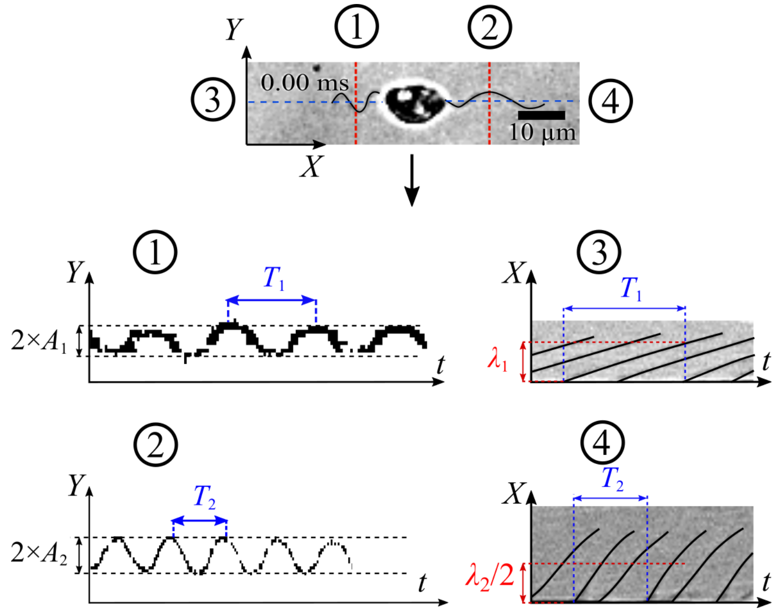


**Fig. S1** Strategy to apply kymograph to obtain characteristics of beating zoospore flagella.

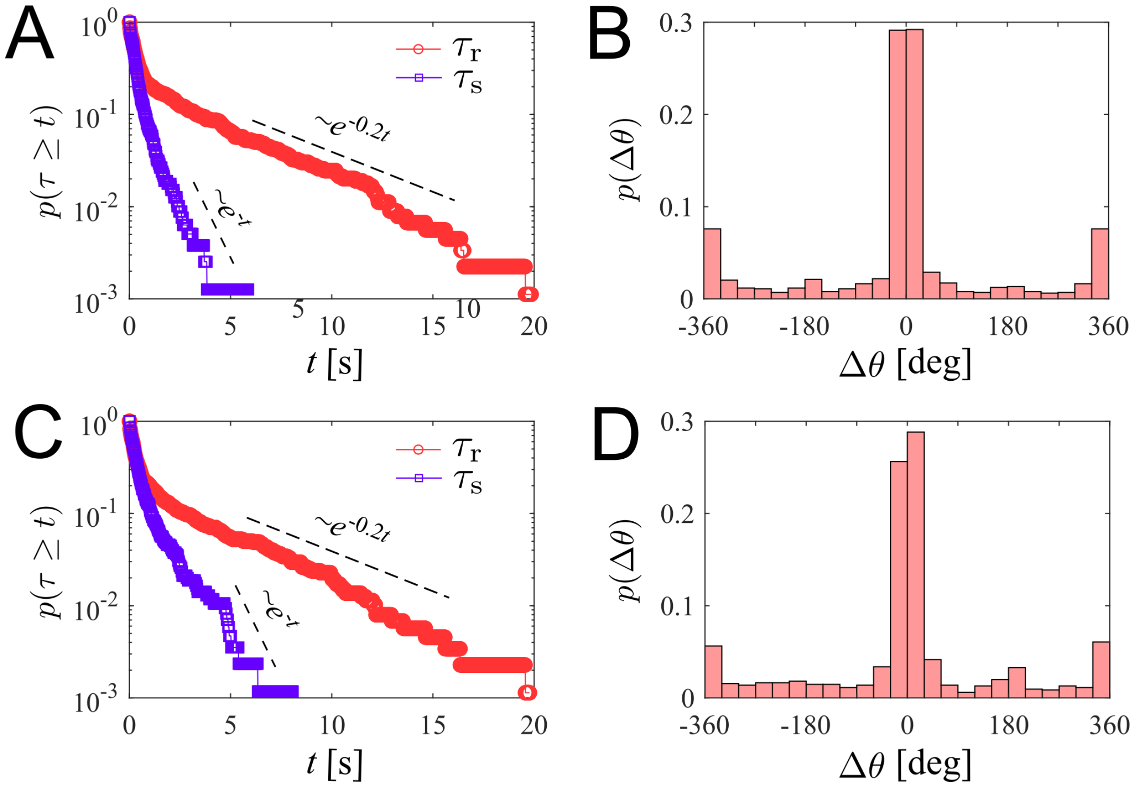


**Fig. S2** Estimation of $\tau_{\rm r}$, $\tau_{\rm s}$ and $\delta\theta$ for different variation of $U_{\rm th}$. (A) $\tau_{\rm r}$, $\tau_{\rm s}$ and (B) $\Delta\theta$ for $U_{\rm th} - 10\,\%$. (C) $\tau_{\rm r}$, $\tau_{\rm s}$ and (D) $\Delta\theta$ for $U_{\rm th} + 10\,\%$

Table 1: Characteristics of helical trajectories of zoospores during straight runs

| Sample | pitch $p$ (µm) | Radius $R$ (µm) | Gyrational speed $\dot{\phi}$ (rads$^{-1}$) |
|---|---|---|---|
| 1 | 122 | 3.250 | 3.64 $\pi$ |
| 2 | 123 | 4.435 | 4.00 $\pi$ |
| 3 | 145 | 4.129 | 2.86 $\pi$ |
| 4 | 164 | 3.390 | 2.5 $\pi$ |
| 5 | 178 | 4.850 | 2.1 $\pi$ |
| 6 | 112 | 3.010 | 4.44 $\pi$ |
| 7 | 120 | 4.300 | 4.44 $\pi$ |
| 8 | 96 | 4.244 | 5 $\pi$ |
| 9 | 127 | 4.090 | 3.33 $\pi$ |
| 10 | 106 | 4.744 | 4 $\pi$ |

### 4 Estimation of mastigoneme density

Mastigoneme density is an important parameter, which determines the thrust reversal effect of the anterior flagellum and generates speed for zoospores. Using TEM, we capture images of anterior flagellum containing the mastigonemes. We first select a longitudinal flagellum length and manually count the total number of mastigonemes attached along that distance (Fig. S3). The mastigoneme density $N_{\mathrm{m}}$ is calculated as the ratio between the number of counted mastigonemes and the selected flagellum length.

Table 2: Measurement data of mastigonemes in TEM images

| Sample | No. of mastigonemes | Flagellum length (µm) | Density (µm$^{-1}$) |
|---|---|---|---|
| 1 | 152 | 11.842 | 12.8357 |
| 2 | 168 | 11.597 | 14.4865 |
| 3 | 113 | 8.627 | 13.0984 |
| 4 | 181 | 13.590 | 13.3186 |
| 5 | 198 | 15.195 | 13.0306 |
| 6 | 177 | 13.994 | 12.6483 |
| 7 | 88 | 7.499 | 11.7349 |

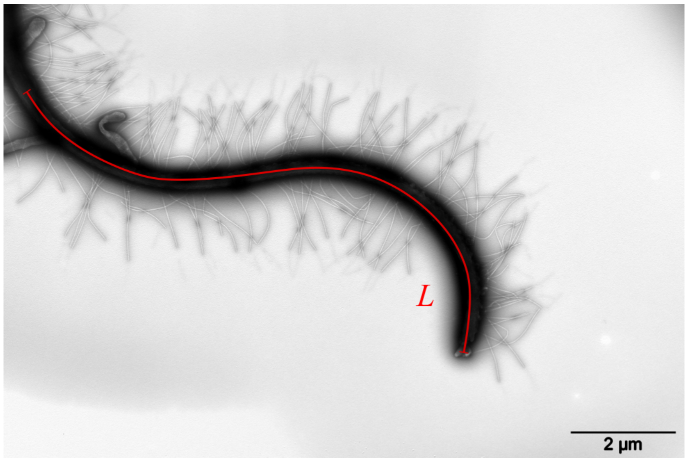


**Fig. S3** Strategy to estimate mastigoneme density.

### 5 Hydrodynamics of two flagella in a swimming zoospore using Resistive Force Theory

Following the schematics of Fig. 3(a) in the main text, we consider a parameter $s_k$ as the distance along the flagellum from the root to the segment $\mathrm{d}s_k$, then we define dimensionless constants $\alpha_k$ as the ratios of the wavelength to the arc length of the anterior ($k = 1$) and posterior flagellum ($k = 2$), respectively. Whereas

$$\alpha_1 = \frac{\lambda_1}{L_1} = \frac{x_1 - c}{s_1} \tag{1}$$

and

$$\alpha_2 = \frac{\lambda_2}{L_2} = \frac{-(x_2 + c)}{s_2}. \tag{2}$$

Thus, we can express the segment position $\vec{r}_k$ in terms of $s_k$, as follows

$$\vec{r}_1 = (\alpha_1 s_1 + c)\vec{i} + A_1 \sin\left(\omega_1 t - \frac{2\pi s_1}{L_1}\right)\vec{j}, \tag{3}$$

and

$$\vec{r}_2 = (-\alpha_2 s_2 - c)\vec{i} + A_2 \sin\left(\omega_2 t - \frac{2\pi s_2}{L_2}\right)\vec{j}. \tag{4}$$

We achieve the velocity of the flagella relative to the cell body as

$$\vec{v}_{\mathrm{f/b}} = d\vec{r}_k/dt = \omega_k A_k \cos\left(\omega_k t - \frac{2\pi s_k}{L_k}\right)\vec{j}. \tag{5}$$

The beating of flagella results in the movement of the cell body relative to water, with a velocity:

$$\vec{v}_{\mathrm{b/w}} = U_X \vec{i}. \tag{6}$$

We neglect the velocity of the cell body in $Y$-direction with the assumption that the zoospore swimming is very directional and $U_X >> U_Y$.

Resistive Force Theory (RFT) states that the drag force by fluid acting on an infinitesimal segment $\mathrm{d}s$ of the flagellum is proportional to the relative velocity of fluid to the flagellum segment, as follows

$$\frac{\mathrm{d}\vec{F}}{\mathrm{d}s} = K_\mathrm{N} V_\mathrm{N} \vec{n} + K_\mathrm{L} V_\mathrm{L} \vec{l}, \tag{7}$$

where $V_\mathrm{N}$ and $V_\mathrm{L}$ are two components of relative velocity of fluid in normal and tangent direction to the flagellum segment, $K_\mathrm{N}$ and $K_\mathrm{L}$ are the drag coefficients of the flagellum in normal and tangent to the flagellum segment, $\vec{n}$ and $\vec{l}$ are the unit vectors normal and tangent to the flagellum segment, respectively.

Equation (7) can be expressed as:

$$\begin{aligned}\frac{\mathrm{d}\vec{F}}{\mathrm{d}s} &= K_\mathrm{N}(\vec{v}_{\mathrm{w/f}} \cdot \vec{n})\vec{n} + K_\mathrm{L}(\vec{v}_{\mathrm{w/f}} \cdot \vec{l})\vec{l} \\ &= K_\mathrm{N}\left(\vec{v}_{\mathrm{w/f}} - (\vec{v}_{\mathrm{w/f}} \cdot \vec{l})\vec{l}\right) + K_\mathrm{L}(\vec{v}_{\mathrm{w/f}} \cdot \vec{l})\vec{l} \\ &= K_\mathrm{N}\left[(\vec{v}_{\mathrm{w/f}} \cdot \vec{l})\left(\frac{K_\mathrm{L}}{K_\mathrm{N}} - 1\right)\vec{l} + \vec{v}_{\mathrm{w/f}}\right],\end{aligned} \tag{8}$$

where $\vec{v}_{\mathrm{w/f}}$ is the relative velocity of water to the flagellum, which is achieved as:

$$\vec{v}_{\mathrm{w/f}} = -\vec{v}_{\mathrm{b/w}} - \vec{v}_{\mathrm{f/b}}. \tag{9}$$

and the tangent unit vector $\vec{l}$ is expressed as:

$$\vec{l} = \frac{1}{|\frac{\partial \vec{r}}{\partial s}|}\frac{\partial \vec{r}}{\partial s}. \tag{10}$$

During the derivative, we use an assumption that the flagella have small amplitude deflection ($A << L$). Thus, we can approximate that $\cos^2\left(\frac{2\pi s_k}{L_k} - \omega_k t\right) \approx 0.5$. This helps to simplify the term $|\frac{\partial \vec{r}}{\partial s}| = \sqrt{\alpha_k^2 + 4\pi^2 \frac{A_k^2}{L_k^2}\cos^2\left(\frac{2\pi s_k}{L_k} - \omega_k t\right)} = \sqrt{\alpha_k^2 + 2\pi^2\frac{A_k^2}{L_k^2}}$.

The drag coefficients $K_N$ and $K_L$ is estimated by Brennen and Winet, as follows:

$$K_{\mathrm{N}} = \frac{4\pi\mu}{\ln\left(\frac{4\lambda}{d}\right) - 2.90}, \tag{11}$$

$$K_{\mathrm{L}} = \frac{2\pi\mu}{\ln\left(\frac{4\lambda}{d}\right) - 1.90}, \tag{12}$$

where $\mu = 8.9 \times 10^{-4}\,\mathrm{Pa\,s}$ is the water viscosity at $25\,^\circ\mathrm{C}$, $\lambda$ is the wavelength and $d$ is the diameter of the flagellum. With $U_Y$ being neglected, we also consider that the two flagella do not generate force in $Y$-direction, but only induce thrust in $X$-direction. Hence,

$$\mathrm{d}\vec{F} \approx \mathrm{d}F_X\vec{i} = (\mathrm{d}\vec{F}\cdot\vec{i})\vec{i}. \tag{13}$$

We then apply RFT on each flagellum of the zoospore to calculate the total drag force acting on it. For the posterior flagellum, each segment $\mathrm{d}s_2$ is a simple smooth and slender filament, having drag coefficients $K_{\mathrm{N2}}$ and $K_{\mathrm{L2}}$. following equation (11) and (12). From equation (8) and (13), we derive the drag force of water acting on posterior flagellum as

$$F_{X,2} = K_{\mathrm{N2}}L_2\left[\frac{-2\pi^2 v_{\mathrm{w2}}(\gamma_2 - 1)\beta_2^2 - (\gamma_2 - 1)U_X}{1 + 2\pi^2\beta_2^2} - U_X\right], \tag{14}$$

where $v_{\mathrm{w2}} = \lambda_2 f_2$ is the wave propagation velocity, $\gamma_2 = K_{\mathrm{L2}}/K_{\mathrm{N2}}$, $\beta_2 = A_2/\lambda_2$. For the anterior flagellum, each segment $\mathrm{d}s_1$ contains additional $N\mathrm{d}s_1$ mastigonemes which are considered perpendicular to the segment itself. These mastigonemes also act as slender filaments experienced drag from water. Interestingly, due to the direction arrangement, the relative velocity normal to the flagellum segment results in drag force in tangent direction to the mastigonemes, and subsequently, the relative velocity tangent to the flagellum segment results in drag force in normal direction to the mastigonemes. Hence, the total drag force acting on a segment of the anterior flagellum is derived as:

$$\frac{\mathrm{d}\vec{F}_1}{\mathrm{d}s_1} = (K_{\mathrm{Nf1}} + NhK_{\mathrm{Lm1}})V_{\mathrm{N1}}\vec{n}_1 + (K_{\mathrm{Lf1}} + NhK_{\mathrm{Nm1}})V_{\mathrm{L1}}\vec{l}_1, \tag{15}$$

where $N$ is the density of mastigonemes, $h$ is the length of each mastigoneme, $V_{\mathrm{N1}}$ and $V_{\mathrm{L1}}$ are two components of relative velocity of fluid in normal and tangent direction to the segment $\mathrm{d}s_1$, $K_{\mathrm{Nf1}}$ and $K_{\mathrm{Lf1}}$ are the drag coefficients in normal and tangent direction of the flagellum filament, respectively; $K_{\mathrm{Nm1}}$ and $K_{\mathrm{Lm1}}$ are the drag coefficients in normal and tangent direction of the mastigonemes, respectively. $K_{\mathrm{Nf1}}$, $K_{\mathrm{Lf1}}$, $K_{\mathrm{Nm1}}$ and $K_{\mathrm{Lm1}}$ are also calculated from equation (11) and (12). In another perspective, we can consider the anterior flagellum receives additional drag from the mastigonemes, which is presented by two increased drag coefficients in normal and tangent direction defined as

$$K_{\mathrm{N1}} = (K_{\mathrm{Nf1}} + N_{\mathrm{m}}hK_{\mathrm{Lm1}}) \tag{16}$$

and

$$K_{\mathrm{L1}} = (K_{\mathrm{Lf1}} + N_{\mathrm{m}}hK_{\mathrm{Nm1}}), \tag{17}$$

respectively. Here, $N_{\rm m}$ is the density of mastigonemes; $h$ is the length of each mastigoneme; $K_{\rm Nf1}$ and $K_{\rm Lf1}$ are the drag coefficients in normal and tangent direction of the flagellum filament, respectively; $K_{\rm Nm1}$ and $K_{\rm Lm1}$ are the drag coefficients in normal and tangent direction of the mastigonemes, respectively. $K_{\rm Nf1}$, $K_{\rm Lf1}$, $K_{\rm Nm1}$ and $K_{\rm Lm1}$ are also calculated from equation (11) and (12). The total fluid drag force acting on the anterior flagellum is also derived from equation 8 and 13:

$$F_{X,1} = K_{\rm N1} L_1 \left[ \frac{2\pi^2 v_{\rm w1}(\gamma_1 - 1)\beta_1^2 - (\gamma_1 - 1)U_X}{1 + 2\pi^2\beta_1^2} - U_X \right], \tag{18}$$

where $v_{\rm w1} = \lambda_1 f_1$ is the wave propagation velocity of the anterior flagellum, $\gamma_1 = K_{\rm L1}/K_{\rm N1}$, $\beta_1 = A_1/\lambda_1$.

At the same time, the ellipsoidal cell body moving with velocity $U_X$ also experiences a drag force from water (following Happel and Brenner)

$$\vec{F}_{d,cell} = -6\pi\mu b \xi_e U_X \vec{i}, \tag{19}$$

where $\mu$ is water viscosity, $\xi_e$ is the shape coefficient of the ellipse cell body in 2D. $\xi_e$ is estimated as

$$\xi_e = \frac{4/3\left(\kappa^2 - 1\right)}{\frac{2\kappa^2-1}{\sqrt{\kappa^2-1}} \ln(\kappa + \sqrt{\kappa^2 - 1}) - \kappa}, \tag{20}$$

with $\kappa = a/b$ being the ellipse body ratio. In low Reynolds number condition, total forces equate to zero due to approximately zero inertia. Thus,

$$\Sigma\vec{F} = \vec{F}_1 + \vec{F}_2 + \vec{F}_{d,cell} = \vec{0}. \tag{21}$$

Combine equation (14), (18) and (19), we achieve translational velocity $U_X$ of the zoospore as shown in Equation 5 of the main texts.

### 6 Power and efficiency of two flagella during in zoospore swimming

We derive the power that each flagellum generates during zoospore swimming to investigate how the two flagella cooperate and contribute to the swimming. The power generated by the anterior flagellum $P_1$ is calculated as

$$P_1 = \int_{s_1=0}^{L_1} \vec{v}_1 \cdot {\rm d}\vec{F}_1, \tag{22}$$

where $\vec{v}_1 = \vec{v}_{\rm w/f(1)}$ is the relative velocity vector of water to each flagellar segment, and ${\rm d}\vec{F}_1$ is the drag force of water acting on each flagellar segment. From (8), we have

$$\vec{v}_1 \cdot {\rm d}\vec{F}_1 = K_{\rm N1} \left[ (\gamma_1 - 1)\left(\vec{v}_1 \cdot \vec{l}_1\right)^2 + \vec{v}_1^2 \right] {\rm d}s_1. \tag{23}$$

Then, we can derive $P_1$ from (9), (10) and (23) as

$$P_1 = K_{\rm N1} L_1 \left[ (\gamma_1 - 1)\frac{\left(2\pi^2 v_{\rm w1}\beta_1^2 - U_X\right)^2}{1 + 2\pi^2\beta_1^2} + U_X^2 + 2\pi^2 v_{\rm w1}^2 \beta_1^2 \right]. \tag{24}$$

Similarly, we can also derive $P_2$ as

$$P_2 = K_{\rm N2} L_2 \left[ (\gamma_2 - 1)\frac{\left(2\pi^2 v_{\rm w2}\beta_2^2 + U_X\right)^2}{1 + 2\pi^2\beta_2^2} + U_X^2 + 2\pi^2 v_{\rm w2}^2 \beta_2^2 \right]. \tag{25}$$

Table S1: Physical parameters of beating flagella of *P. parasitica* zoospores in normal conditions.

| **Structures** | **Parameters** | **Denotes** | **Value** |
|---|---|---|---|
| Anterior flagellum | Amplitude (μm) | $A_1$ | 1.6 |
| | Wavelength (μm) | $\lambda_1$ | 7.5 |
| | Length (μm) | $L_1$ | 23.4 |
| | Beating freq. (Hz) | $f_1$ | 74 |
| | Drag coeff. of filaments | $K_{\mathrm{Nf1}}$ | $6.5589 \cdot 10^{-15}$ |
| | ($\mathrm{N\,s\,\mu m^{-2}}$) | $K_{\mathrm{Lf1}}$ | $2.067 \cdot 10^{-15}$ |
| | Drag coeff. of mastigonemes | $K_{\mathrm{Nm}}$ | $2.7906 \cdot 10^{-15}$ |
| | ($\mathrm{N\,s\,\mu m^{-2}}$) | $K_{\mathrm{Lm}}$ | $1.1167 \cdot 10^{-15}$ |
| | Mastigoneme length (μm) | $h$ | 1.5 |
| | Mastigoneme density ($\mathrm{\mu m^{-1}}$) | $N_{\mathrm{m}}$ | 15 |
| | ($A_1/\lambda_1$) ratio | $\beta_1$ | 0.21 |
| | Drag coeff. ratio ($K_{\mathrm{L1}}/K_{\mathrm{N1}}$) | $\gamma_1$ | 2.047 |
| Posterior flagellum | Amplitude (μm) | $A_2$ | 2.9 |
| | Wavelength (μm) | $\lambda_2$ | 15.17 |
| | Beating freq. (Hz) | $f_2$ | 128 |
| | Length (μm) | $L_2$ | 25.8 |
| | Drag coefficients | $K_{\mathrm{N2}}$ | $4.6415 \cdot 10^{-15}$ |
| | ($\mathrm{N\,s\,\mu m^{-2}}$) | $K_{\mathrm{L2}}$ | $1.6401 \cdot 10^{-15}$ |
| | ($A_2/\lambda_2$) ratio | $\beta_2$ | 0.19 |
| | Drag coeff. ratio ($K_{\mathrm{L2}}/K_{\mathrm{N2}}$) | $\gamma_2$ | 0.3534 |

The useful power $P_0$ required to propel the cell body with a speed $U_X$ can be derived as

$$P_0 = F_{d,cell} \cdot U_X = 6\pi\mu b\xi_e U_X^2. \tag{26}$$

The efficiency of the two flagella in propelling the cell body is estimated as

$$\eta = \frac{P_0}{P_1 + P_2}. \tag{27}$$

### 7 Dimensions and physical parameters of zoospore's flagella

With the characteristics of beating flagella obtained by kymograph, we calculate the values of drag coefficients $K_{\mathrm{N}}$, $K_{\mathrm{L}}$, flagellum shape ratio $\beta$, and drag coefficient ratio $\gamma$. The results are presented in Table S1.

### 8 Supplementary Movies

**Supp. Movie 1.** Zoospores swimming in water. The movie was captured in the microscopic assay, with objectives 4×, 60 fps, duration 60 s. The trajectories of zoospores were tracked by combining TrackMate (in Fiji) and manual tracking.
**Supp. Movie 2.** A zoospore swimming near water/air interface.
**Supp. Movie 3.** The result from our simulation with the strategy described in Fig. 2(h). The parameters of the simulation are extracted from the statistics of the swimming of zoospores, presented in Fig. 2(d-g).

**Supp. Movie 4.** An individual zoospore swimming in water. It is observed to swim straight with the helical trajectory due to the body self-rotation, then perform a 180° turning.
**Supp. Movie 5.** A zoospore performing a turning event, which involves reducing speed, body rotation and steering resulting from active gait changing of anterior flagellum. The posterior flagellum is immobile during the turning event.
**Supp. Movie 6.** A temporal zoom to the "Rotate" step of the turning event, showing a zoospore changing gait from sinuisoidal wave to power and recovery stroke similarly to *C. reinhardtii*'s.

### 9 Data publication

Our experimental data are accessible via Zenodo (URL: https://doi.org/10.5281/zenodo.4710633). In the data, we include

(1) datasets of all zoospore positions along multiple trajectories in the experiment of Figure 2,

(2) a MATLAB file to compute all the statistical results in Figure 2(d-g),

(3) a MATLAB file containing the simulation model presented in Figure 2(h), and

(4) datasets of zoospore positions, speed, moving directions, body orientations during the turning, presented in Figure 4.

During the review process, the data are currently restricted. Only reviewers can access the file using a password. The password will be removed afterwards.
